# Neural signatures of predictive language processing in parkinson’s disease with and without mild cognitive impairment

**DOI:** 10.1101/2020.11.23.392647

**Authors:** Patricia León-Cabrera, Javier Pagonabarraga, Joaquín Morís, Saúl Martínez-Horta, Juan Marín-Lahoz, Andrea Horta-Barba, Helena Bejr-Kasem, Jaime Kulisevsky, Antoni Rodríguez-Fornells

## Abstract

Cognitive deficits are common in Parkinson’s disease (PD), with some PD patients meeting criteria for mild cognitive impairment (MCI). An unaddressed question is whether linguistic prediction is preserved in PD. This ability is nowadays deemed crucial in achieving fast and efficient comprehension, and it may be negatively impacted by cognitive deterioration. To fill this gap of knowledge, we used event-related potentials (ERPs) to evaluate mechanisms of linguistic prediction in a sample of PD patients (on dopamine compensation) with and without MCI. To this end, participants read sentence contexts that were predictive or not about a sentence-final word. The final word appeared after 1 second, matching or mismatching the prediction. The introduction of the interval allowed to capture neural responses both before and after sentence-final words, reflecting semantic anticipation and processing. PD patients with normal cognition (N = 58) showed ERP responses comparable to those of matched controls. Specifically, in predictive contexts, a slow negative potential developed prior to sentence-final words, reflecting semantic anticipation. Later, expected words elicited reduced N400 responses (compared to unexpected words), indicating facilitated semantic processing. Besides, PD patients with MCI (N = 20) showed a prolongation of the N400 congruency effect (compared to matched PD patients without MCI), indicating that further cognitive decline impacts semantic processing. Finally, lower verbal fluency scores correlated with prolonged N400 congruency effects and with reduced pre-word differences in all PD patients (N = 78). This relevantly points to a role of deficits in temporal-dependent mechanisms in PD, besides prototypical frontal dysfunction, in altered semantic anticipation and semantic processing during sentence comprehension.

## 1. INTRODUCTION

Parkinson’s disease (PD) is a chronic, neurodegenerative disorder that, in addition to motor defects, involves difficulties in a variety of cognitive domains (Kudlicka *et al*., 2011; Muslimovic *et al*., 2005). Patients with PD may exhibit significant problems with language comprehension and language production in everyday life. These difficulties have been partly explained by studies that explored in-depth linguistic function in sentence comprehension (for a review, see Pell *et al*., 2008) and have been mostly attributed to syntactic alterations (Lieberman, 1992; Friederici *et al*., 2002), although there is also evidence pointing to slower or delayed lexical and semantic activation (Arnott *et al*., 2001; Angwin *et al*., 2005; Angwin *et al*., 2017). Moreover, cognitive resource limitations in functions that enable or support language processing, such as executive functions or working memory, may secondarily impact comprehension (Grossman *et al*., 2002). Yet, there is still paucity of data in the literature on language processing and associated cognitive disturbances in non-demented PD patients.

An aspect of language comprehension that has not been evaluated in PD is predictive language processing, which plays an important role in achieving fast and efficient language processing (Kutas *et al*., 2011). Accordingly, readers and listeners probabilistically infer and pre-activate different aspects of upcoming words (e.g., van Berkum *et al*., 2005; Wicha, *et al*., 2003). Much evidence of language prediction has been derived from the N400 event-related potential (ERP) component, an index of semantic processing (Kutas *et al*., 1980). The N400, a negativity peaking ~400 ms after word onset, is reduced for words that are more predictable in a given context (N400 context effect) (Kutas *et al*., 1984), which is considered to reflect facilitated semantic processing owing to the prediction of the word or some of its semantic features (Federmeier *et al*., 1999). Most relevantly, recent research has shown that processing differences arise even before the word is presented. In particular, slow negative potentials (SNP) consistent with semantic anticipation precede sentence-final words in predictive contexts (e.g., before “ball” in “The goalkeeper managed to catch the… ball”) (León-Cabrera *et al*., 2017; 2019; Grisoni *et al*., 2017; for a review, see Pullvermüller *et al*., 2020).

Importantly, prior research suggests that linguistic prediction is reduced in populations with limited cognitive resources (Federmeier *et al*., 2002; 2005; 2010; Payne *et al*., 2008; Wlotko *et al*., 2012; for reviews, see Payne *et al*., 2019; Huettig, 2015). For instance, older adults seem to take less advantage of semantically rich contexts to facilitate subsequent processing (Federmeier *et al*., 2005) and those with lower verbal fluency scores exhibit diminished or absent ERP effects associated with prediction, suggesting the adoption of a ‘wait-and-see’, incremental strategy instead (Federmeier *et al*., 2002). Crucially, up to 40% of individuals with PD will develop mild cognitive impairment (PD-MCI) within the first 5 years of disease (Aarsland *et al*., 2010; Litvan *et al*., 2011), which is a robust predictor of further conversion to dementia (PDD). Among the several cognitive phenotypes characterizing PD-MCI, changes in visuoperceptive skills, memory or language have been highlighted to be good predictors of the conversion from PD-MCI to PDD (Horta-Barba *et al*., 2020; Martínez-Horta *et al*., 2019; Lang *et al*., 2019). Based on this, we hypothesized that deficits in predictive language processing might contribute to sentence comprehension difficulties in PD and in PD-MCI.

To address this, the current study investigated ERP signatures of semantic prediction and semantic processing in PD with normal cognition (PD-NC) and PD-MCI. We presented sentence contexts that were predictive or not of a sentence-final word that was delayed by 1 s (see **Table 2**) (León-Cabrera *et al*., 2019), thus allowing to capture the abovementioned ERP signatures in the anticipatory and processing stages of prediction. In predictive contexts, half of the final words were semantically incongruent, thus representing a prediction mismatch. Lastly, following prior findings in older adults (Federmeier *et al*., 2002), we explored whether the targeted ERP signatures of prediction were reduced or diminished in PD patients with lower verbal fluency scores.

## 2. MATERIALS AND METHODS

### 2.1. Participants

Participants were prospectively recruited from a sample of outpatients regularly attending the Movement Disorders Clinic at Hospital de la Santa Creu i Sant Pau (Barcelona, Spain). They were invited to participate in a longitudinal study involving exhaustive neuropsychological testing and two EEG recording sessions (one at baseline and a follow-up after 1 year). Healthy adults were also invited to participate in the study to serve as controls. The project was approved by the research ethics committee of the Hospital de la Santa Creu i Sant Pau. Before inclusion, written informed consent was obtained from all the participants.

### 2.2. Clinical and cognitive testing

PD patients had been diagnosed by a neurologist with expertise in movement disorders. All patients accomplished steps 1 and 2 of the UKPDSBB criteria, and three or more of the four first supportive positive criteria of step 3 (Hughes *et al*., 1992). Motor status and stage of illness were assessed by the MDS-UPDRS-III and Hoehn & Yahr scales (Goetz *et al*., 2008). Demographic variables including age, gender, years of formal education, disease onset, medication history, as well as the levodopa equivalent daily dosage (LDD) (Tomlinson *et al*., 2010) were collected for all patients (**Table 1**). All participants were classified as having either normal cognition or mild cognitive impairment (MCI) based on their score in the Parkinson’s Disease - Cognitive Rating Scale (PD-CRS) (Pagonabarraga *et al*. 2008). A cutoff score of ≤ 83 was used to classify patients as with or without PD-MCI (Fernández de Bobadilla *et al*. 2013). Participants with dementia were excluded according to the MDS diagnostic criteria for PDD (Emre *et al*., 2007). In PD patients, cognition was examined during the ‘on’ state, and all participants were on stable doses of dopaminergic drugs during the 4 weeks before inclusion. In addition, their semantic and phonological verbal fluency was evaluated. For each participant, total verbal fluency scores were obtained by averaging the direct scores in semantic and phonological fluency subtests.

**Table 1.** Demographics and clinical features of the patient and control samples.

|  | CG | PD-NC | P | sPD-NC | PD-MCI | P |
| --- | --- | --- | --- | --- | --- | --- |
| N | 24 | 58 |  | 19 | 20 |  |
| Age (years) | 62.85<br>(6.82)<br>[59.9, 65.7] | 64.47<br>(7.61)<br>[62.4, 66.4] | .37 | 70.75<br>(2.70)<br>[69.4, 72] | 72.22<br>(4.48)<br>[70.1, 74.3] | .22 |
| Education (years) | 14.5<br>(4.04)<br>[12.7, 16.2] | 13.2<br>(4.57)<br>[12, 14.4] | .23 | 12.42<br>(5.08)<br>[9.9, 14.8] | 10.4<br>(5.16)<br>[7.9, 12.8] | .22 |
| Women (%) | 41.6 | 31 | .35 | 42.1 | 45 | .85 |
| Disease duration (years) | n/a | 5.26<br>(3.00)<br>[4.4, 6] | - | 5.68<br>(3.06)<br>[4.2, 7.1] | 6.28<br>(3.8)<br>[4.4, 8] | .59 |
| Levodopa dose (mg/d) | n/a | 265.25<br>(249.58) | - | 252.87<br>(293.74) | 384.4<br>(215.04) | .12 |
| LEDD (mg/d) | n/a | 456.04<br>(230.68) | - | 397.18<br>(236.08) | 532.83<br>(209.70) | .09 |
| PD-CRS total score [$\leq 83$ cutoff] | 106.16<br>(8.74)<br>[102.4, 109.85] | 99.72<br>(10.86)<br>[96.8, 102.5] | .01 | 98.2<br>(9.17)<br>[93.7, 102.6] | 67.15<br>(5.39)<br>[64.6, 69.6] | <.001 |
| Total verbal fluency (DS) | 19.66 (3.12) | 18.01 (4.18) | .08 | 17.63 (4.32) | 10.52 (2.08) | <.001 |

The values correspond to means, standard deviations (in round brackets) and 95% mean confidence intervals (CI) (in square brackets). For simplicity, CI are shown only for the relevant variables in the group comparisons.
P-values were determined with independent samples t-tests between groups for continuous and ordinal variables and with Pearson Chi-Square for nominal variables (Gender).
n/a = not applicable; LEDD = Total levodopa equivalent daily dosage; PD-CRS = Parkinson's Disease Cognitive Rating Scale; DS = Direct score.

### 2.3. Final samples

A total of 135 participants completed the EEG recording session. Of those participants, 102 were PD patients and 33 were healthy controls. The data of some participants were excluded because the files were corrupt (n = 3) or their recordings were excessively noisy (n = 4) as determined by visual inspection of the raw data, leaving 128 participants available for analyses. From the remaining participants, 97 were PD patients and 31 were controls. Controls were excluded if they had mild cognitive impairment (n = 3) or their cognitive status was not available (n = 2). Lastly, participants with less than 20 available trials in any condition after EEG preprocessing (n = 9) were also excluded from the analyses (see EEG preprocessing section for more information about the artefact detection and rejection procedure).

The final pool of data consisted of 88 PD patients and 26 controls, from which matched samples were subsequently obtained to perform the following comparisons according to the goals of the present study: 1) PD patients with normal cognition (PD-NC) (N = 58) with an age-, gender- and education-matched Control Group (CG) (N = 24), and 2) a subsample of Parkinson’s disease patients with normal cognition (sPD-NC) (N = 19) with an age-, gender-, education- and years of disease’s evolution-matched PD patients with mild cognitive impairment (PD-MCI) (N = 20). To avoid the comparison of patients classified as sPD-NC or sPD-MCI but having similar PD-CRS total score, we selected participants with PD-CRS scores of < 75 for the sPD-MCI group and of > 85 for the sPD-NC.

The demographic and clinical features of the final samples are reported in Table 1. CG and PD did not differ in age (t(1,80) = −0.90, *p* = .37, 95% CI [−5.19, 1.95]), education (t(1,80) = 1.20, *p* = .23, 95% CI [−0.84, 3.43]), gender (*X*^2^(1, 82) = .853, *p* =.35), or verbal fluency scores (t(1,80) = 1.73, *p* = .08, 95% CI [−0.23, 3.53]). They differed in PD-CRS total scores (t(1,80) = 12.96, *p* <.001, 95% CI [26.2, 35.91]), but both groups were above the established score cutoff of 83 for PD-MCI diagnosis. Likewise, sPD-NC and sPD-MCI did not differ in age (t(1,37) = −1.23, *p* = 0.22, 95% CI [−3.89, .94]), education (t(1,37) = −1.30, *p* = 0.22, 95% CI [−1.30, 5.34]), gender (*X*^2^(1, 39) = .033, *p* =.855), disease’s duration (t(1,37) = −0.53, *p* = .59, 95% CI [−2.84, 1.65]) or levodopa equivalent daily dose (t(1,37) = −1.74, *p* = .09, 95% CI [−293.94, 22.64]), but differed in PD-CRS total score (t(1,37) = 12.96, *p* <.001, 95% CI [26.2, 35.91]) and verbal fluency scores (t(1,37) = 6.59, *p* <.001, 95% CI [4.92, 9.29]). For all comparisons, equal variances were assumed based on null results in Levene’s test.

### 2.4. Materials

We used the materials from León-Cabrera *et al*. (2019) consisting of high (HC) or low (LC) constraining sentence contexts, establishing a stronger or weaker expectation for an upcoming word (see Table 2 for sentence examples). A total of 312 sentences were included, of which 208 had HC sentence contexts (66,6 %) and 104 had LC sentence contexts (33,3 %). The sentences were originally created and categorized as either HC (mean cloze probability = 76%, SD ± 17.7%) or LC (mean cloze probability = 6.1%, SD ± 10.3%) (Mestres-Missé *et al*., 2007). Within the HC condition, half of the sentences ended with a congruent word and half with a semantically incongruent word. For the sake of task brevity to avoid fatigue, we decided to include this contrast (congruent vs. incongruent contrast) only in the HC condition. Sentences were further divided in two lists of 156 sentences, each list containing half of the sentences of each condition, and the use of one or another list was pseudorandomized across participants. In sum, each participant read the following sentences: 52 HC with congruent endings (HC congruent), 52 HC with incongruent endings (HC incongruent), and 52 LC with congruent endings (LC congruent) (**Table 2**). All final words were nouns. Congruent ones were the best completions – the word with the highest cloze-probability (i.e. percentage of individuals that supply that word as a continuation for that sentence) (Taylor, 1953) – and incongruent words were selected from the ESPAL database (Duchon *et al*., 2013) so that they matched the congruent endings in the variables of mean word length, mean number of syllables, word frequency, familiarity, imaginability and concreteness.

**Table 2.**
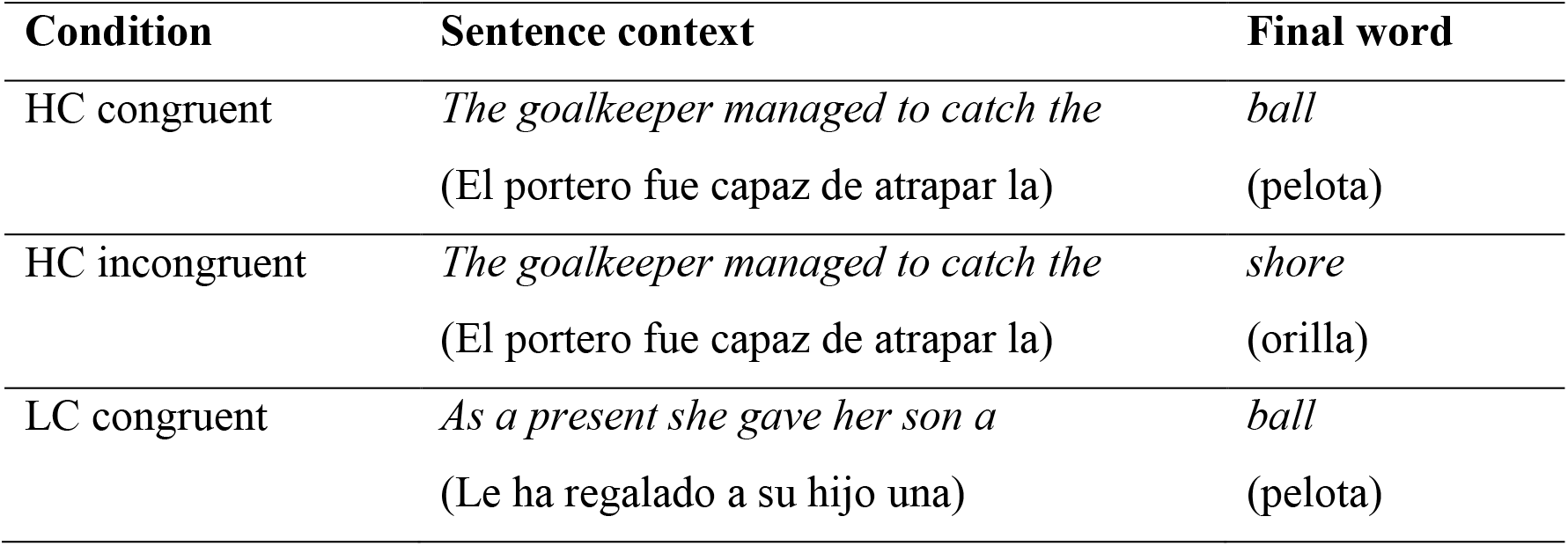
Example sentences of each condition. (original sentences in Spanish in brackets).

### 2.5. Procedure

Participants were tested individually. After external electrodes and the EEG cap were placed, electrode impedances were checked. Participants were instructed to keep fixation on the center of the screen and to blink only during the blinking interval. After completing a short training block (4 sentences), the proper experiment started. They were instructed to read attentively and silently the words that would appear at the center of the screen.

The experiment consisted of 26 blocks of 6 trials each (2 HC congruent, 2 HC incongruent, 2 LC congruent). The order of the sentences within each block was randomized. Words were white on black background (font type: Arial; font size: 24 points). On each trial, a fixation point (a cross) appeared at the center of the screen (800 ms). Then, a sentence was presented word-by-word (300 ms per word, followed by a 200 ms blank screen). Between the penultimate and the final word, the blank screen remained for a period of 1 s (pre-word interval). After the final word (300 ms duration), a blank screen was presented (1000 ms) before the blinking signal (1 s; depiction of an eye at the center of the screen). The next trial began after the offset of the blinking signal (see **Figure 1** for a simplified display of the trial structure).

**Figure 1.**
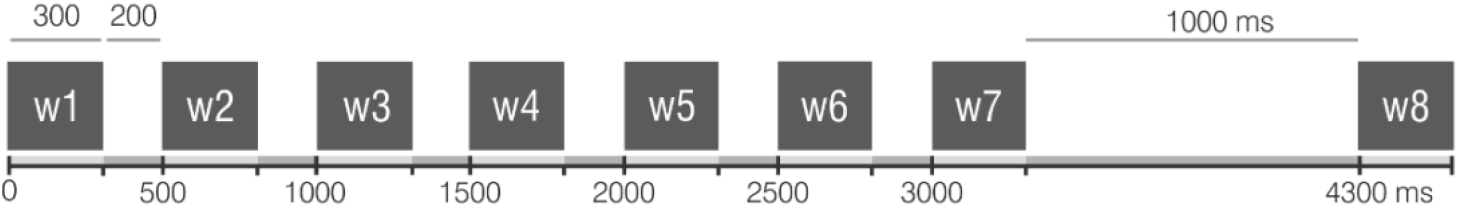
Depiction of the structure of a trial. Sentences were presented one word a time. Each word of the sentence context (w1 to w7) was displayed for 300 ms followed by a 200 ms inter-word interval, except for the last word (w7), which was followed by a 1000 ms interval (pre-word interval) leading up to the presentation of the final word (w8).

To ensure that participants were reading attentively, after every 2 blocks they were asked to judge whether the previous sentence was congruent or not (“did the previous sentence make sense?”). The question remained on the screen until they answered (yes/no) using the keyboard. The next block started immediately after the test except for even-numbered blocks, which were followed by a resting period that could be resumed anytime by the participant.

### 2.6. EEG recording

#### 2.6.1. EEG acquisition

Continuous electroencephalogram (EEG; sampling rate = 250 Hz) was recorded from 19 electrodes (FP1/2, F3/4, F7/8, Fz, C3/4, Cz, P3/4, Pz, T3/4, T5/6, O1/2) mounted on a cap following the international 10−20 system positions. The EEG signal was amplified on-line (band-pass filter = 0.016 – 1000 Hz) with Brain Vision. To monitor ocular artefacts, electrooculogram (EOG) electrodes were recorded with electrodes at dipolar vertical (supraorbital and suborbital ridge of the right eye) and horizontal (external ocular canthus of the left and right eyes) placements. All electrode impedances were kept below 5 kΩ. An on-line notch filter (50 Hz) was applied to attenuate high-frequency electrical noise and data were referenced to the mean activity of the left and right mastoid electrodes.

#### 2.6.2. EEG preprocessing

EEG data were preprocessed using ERPlab v7.0.0 (López-Calderón & Luck, 2014) of the EEGlab toolbox v14.0.0 (Delorme *et al*., 2004), FieldTrip version 20181231 (Oostenweld *et al*., 2011) and custom scripts programmed in MATLAB R2017b. For each participant and condition, the continuous data were segmented into the following epochs of interest: 1) from −100 to 1300 ms from the onset of the penultimate word (pre-word interval), and 2) from −100 to 1000 ms from the target word onset (word interval). Artefact detection and rejection were applied on the epoched data. To facilitate artefact rejection, we computed the vertical (vEOG) and horizontal (hEOG) bipolar EOG channels by subtracting the inferior from the superior vertical EOG channels and the right from the left horizontal EOG channels, respectively. For the pre-word interval, all epochs with activity ± 85 μV in the ocular channels or ± 200 μV in any other channel were automatically removed and the remaining epochs were visually inspected and excluded if they contained blinks, muscle activity or large drifts. For the word interval, the concatenated epochs were first subjected to independent component analysis (ICA) decomposition to correct blinks because preliminary visual inspection revealed that many participants had difficulties refraining from blinking until the blinking signal, leading to a large loss of trials in this interval. ICA was run using the “runica” algorithm as implemented in EEGlab toolbox v14.0.0, excluding the ocular channels from the decomposition. The ocular components were detected and rejected based on visual inspection. After artifact-correction, epochs with activity ± 200 μV in any but ocular channels were automatically rejected, and those that remained were visually screened for any previously unnoticed artefacts (e.g. muscle activity or large drifts).

For the CG with PD-NC comparison, an average of 89.8 % trials per condition were available for analysis (HC = 42.8 trials; LC = 41.9 trials; HCC = 49.6 trials; HCI = 49.5 trials; LCC = 49.7 trials; overall average = 46 trials per condition), with no significant differences in the percentage of available trials between groups (all p-values > 0.3). For the sPD-NC with sPD-MCI comparison, an average of 87.4% trials per condition were included (HC = 39.5 trials; LC = 39.2 trials; HCC = 49.3 trials; HCI = 49.4 trials; LCC = 49.5 trials; overall average = 45 trials per condition), with no significant differences in the percentage of remaining trials between groups (all p-values > 0.1).

### 2.7. Statistical analyses

#### 2.7.1. Event-related potential analyses

Before performing statistical analyses, the data was baseline-corrected to the 100 ms pre-stimulus period and data were low-pass filtered (as implemented in ERPlab v7.0.0). For the pre-word interval, a 5 Hz low-pass Butterworth filter (12 dB/oct roll-off) was applied to the data to focus exclusively on slow activity (Brunia *et al*., 2011). For the word interval, in which the N400 component was targeted, a 30 Hz low-pass Butterworth filter (12 dB/oct roll-off) was used. For the figures presented, the data were filtered using a 12 Hz low-pass filter (roll-off of 12 dB/oct).

#### 2.7.2. Cluster-based permutation analyses

Non-parametrical cluster-based permutation tests were used to assess the differences (Maris and Oostenveld, 2007) (as implemented in Fieldtrip version 20181231 under MATLAB R2017b), a method that exerts proper control for the increased probability of false positives in the context of many comparisons. Broadly, it identifies adjacent time points and channels with similar differences between conditions. The test worked as follows in the present study. For within-subject comparisons, every sample (channel x time) was compared between two conditions (e.g., HC vs. LC) through a dependent *t*-test. For between-subject comparisons, an independent *t*-test compared the effect (e.g., HC minus LC) between groups at each sample (channel x time). Next, adjacent samples were clustered based on a *t*-value threshold (pre-determined from a one-tailed *t*-distribution with an alpha level of .05 and N-1 degrees of freedom) and the cluster with the largest sum of t-values was selected (cluster-level *t*-value). To determine whether effects were significant, the Monte Carlo method was used to construct a null distribution from the cluster-level *t*-values of random partitions obtained by randomly swapping the samples (between conditions and within participants for within-subject comparisons; and between groups and within participants for between-subject comparisons) (5000 randomizations). Only observed clusters with cluster-level *t*-values within the 2.5th percentiles (alpha level of .05) of the null distribution were considered significant.

Choosing one-tailed testing is an optimal methodological decision when there are a priori hypotheses about the direction of the differences (Lakens, 2016). It allows to improve statistical sensitivity, which is particularly important when working with clinical samples. In this case, we expected more negative amplitudes for HC than LC in the pre-word interval (León-Cabrera *et al*., 2017; 2019). As for the interval after the presentation of the word, in which N400 context effects were targeted, we expected more negative amplitudes (larger N400 response) for unexpected that for expected words, that is, for HCI than for HCC, as well as for LCC relative to HCC (e.g., Kutas *et al*., 2011).

#### 2.7.3. Correlations between ERP effects and verbal fluency

After examining ERP effects, we conducted non-parametrical Spearman-Brown correlations to evaluate a potential linear relationship between the ERP effects and verbal fluency performance. All PD patients were included in this analysis (N = 78), that is, PD-NC (N = 58) and PD-MCI (N = 20). For each participant, the total verbal fluency score was obtained by averaging the direct scores in semantic and phonological fluency subtests (see **Supplementary Materials** for correlations with each subtest individually). ERP measures were labeled and quantified as follows: 1) *SNP* (LC minus HC mean amplitude difference in the 600 ms before pre-word onset at a left anterior cluster including electrodes FP1, F3, Fz, F7); 2) *N400 congruency effect* (HCC minus HCI mean amplitude difference from 300 to 500 ms post-word onset); 3) *N400 constraint effect* (LCC minus HCC mean amplitude difference from 300 to 500 ms post-word onset); and, based on the ERP results, 4) *prolonged N400 congruency effect* (HCC minus HCI mean amplitude difference from 600 to 800 ms post-word onset). All effects tied to the N400 were tested on a centrally distributed cluster (averaged electrodes: Cz, C3, C4, Pz, P3, P4). All values were normalized before performing correlations. All *p*-values were corrected with the Holm-Bonferroni correction (Holm, 1979).

## 3. RESULTS

### 3.1. Event-related potentials

#### 3.1.1. CG and PD-NC

**Figures 2A** and **2B** show the grand-average ERPs at the F7 electrode during the pre-word interval in the CG group and the PD-NC group, respectively. The ERPs were computed in the interval from the onset of the penultimate word to the onset of the final word (1300 ms duration, the first 300 ms correspond to the penultimate word, and the last 1000 ms to the delay interval), with a 100 ms pre-stimulus baseline.

**Figure 2.**
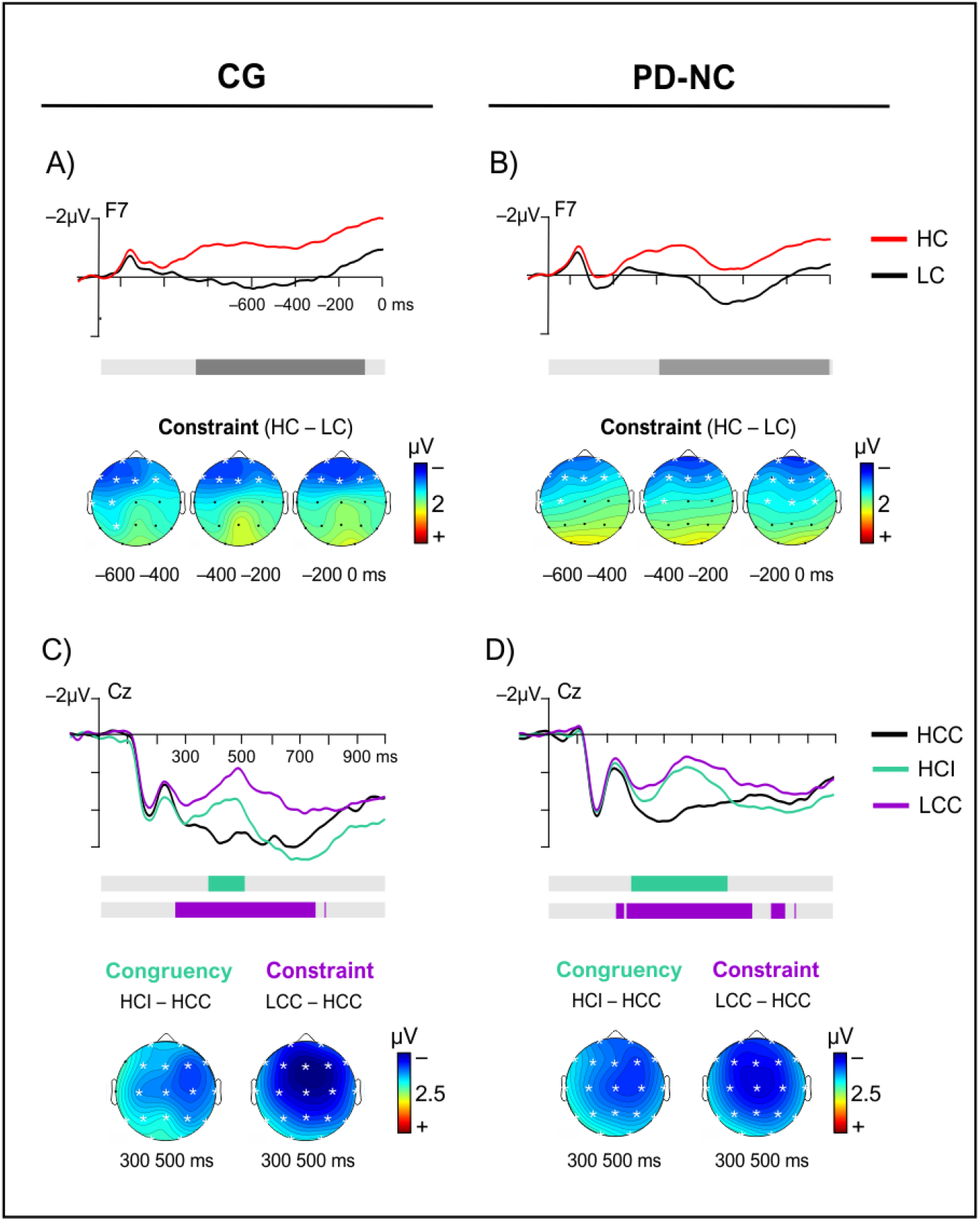
ERP results for CG and PD groups. **A and B)** Grand-averaged ERPs at the F7 electrode during the pre-word interval for both conditions for (A) CG and (B) PD. ERPs are time-locked to the presentation of the penultimate word (100 pre-stimulus baseline). The horizontal bar highlights (grey) the time interval in which significant differences were obtained. Below the ERPs, topographical maps indicating the electrodes (white asterisks) that showed a significant effect of constraint (HC minus LC) in the 600 ms prior to the critical word (in steps of 200 ms). **C and D)** Grand-averaged ERPs at Cz electrode after the presentation of the sentence-final word (100 pre-stimulus baseline) for (C) CG and (D) PD. The horizontal bars highlight the time interval of the N400 effect of Congruency (green) and Constraint (purple). Topographical maps with electrodes (white asterisks) that showed effects in the time interval in which N400 effects were expected to be maximal (300 to 500 ms post-word onset).

Both groups exhibited similar patterns, namely, an N100 associated with the processing of the penultimate word, followed by a positivity, and the later development of a negativity that became progressively larger as the presentation of the final word approached. As expected, based on previous findings in healthy adult population (León-Cabrera *et al*., 2019), the negativity was more prominent in the HC condition (relative to the LC condition), that is, when the final word could be strongly predicted from the prior context. This difference was confirmed by a significant cluster in the CG group (p = 0.01) (**Fig. 2A**) as well as in the PD-NC group (p = 0.01) (**Fig. 2B**). The cluster showed a similar temporal profile in both groups. It started approximately about 800 ms before the final word and went on until the word appeared. In regard to the spatial distribution, the cluster encompassed mainly frontal electrodes, with a slight left lateralization. There were no significant differences between groups.

We then turned to the interval after word presentation. As can be observed in **Figures 2C** and **2D**, for both groups, unexpected words (HCI and LCC) elicited a larger negativity peaking about 400 to 600 ms (relative to expected words; HCC) that is consistent with the canonical features of the N400. These differences were reflected in the following significant clusters.

The effect of constraint (HCC versus LCC) was captured by significant clusters in the CG group (p = .001) (**Fig. 2C**) and in the PD-NC group (p <.001) (**Fig. 2D**), whereby words presented in less predictive contexts (LCC) showed more negative amplitudes than words in strongly predictive contexts (HCC). There were no differences between groups. The cluster started about 250-300 ms after word onset and resolved at 700 ms, and it exhibited a widespread distribution over the scalp. In fact, the cluster was significant at all sites. On the other hand, the effect of congruency (HCC versus HCI) was reflected in a shorter-lived cluster in the CG group (p = .007) (**Fig. 2C**) and PD-NC group (p<.001) (**Fig. 2D**), in this case, confirming more negative amplitudes for incongruent than congruent words in strongly predictive contexts. Again, there were no significant differences between groups. The effect was widespread with a seemingly slight rightward focus in magnitude.

#### 3.1.2. sPD-NC and PD-MCI

The mean voltage pattern elicited during the pre-word interval (**Fig. 3A** and **Fig. 3B**) exhibited the normative N100 component, followed by a transient positivity and a subsequent negativity in the last part of the interval that evidenced the hypothesized tendency towards more negative amplitudes for HC than LC. However, in this case, there were no statistically significant differences between conditions, nor any differences between the two groups.

**Figure 3.**
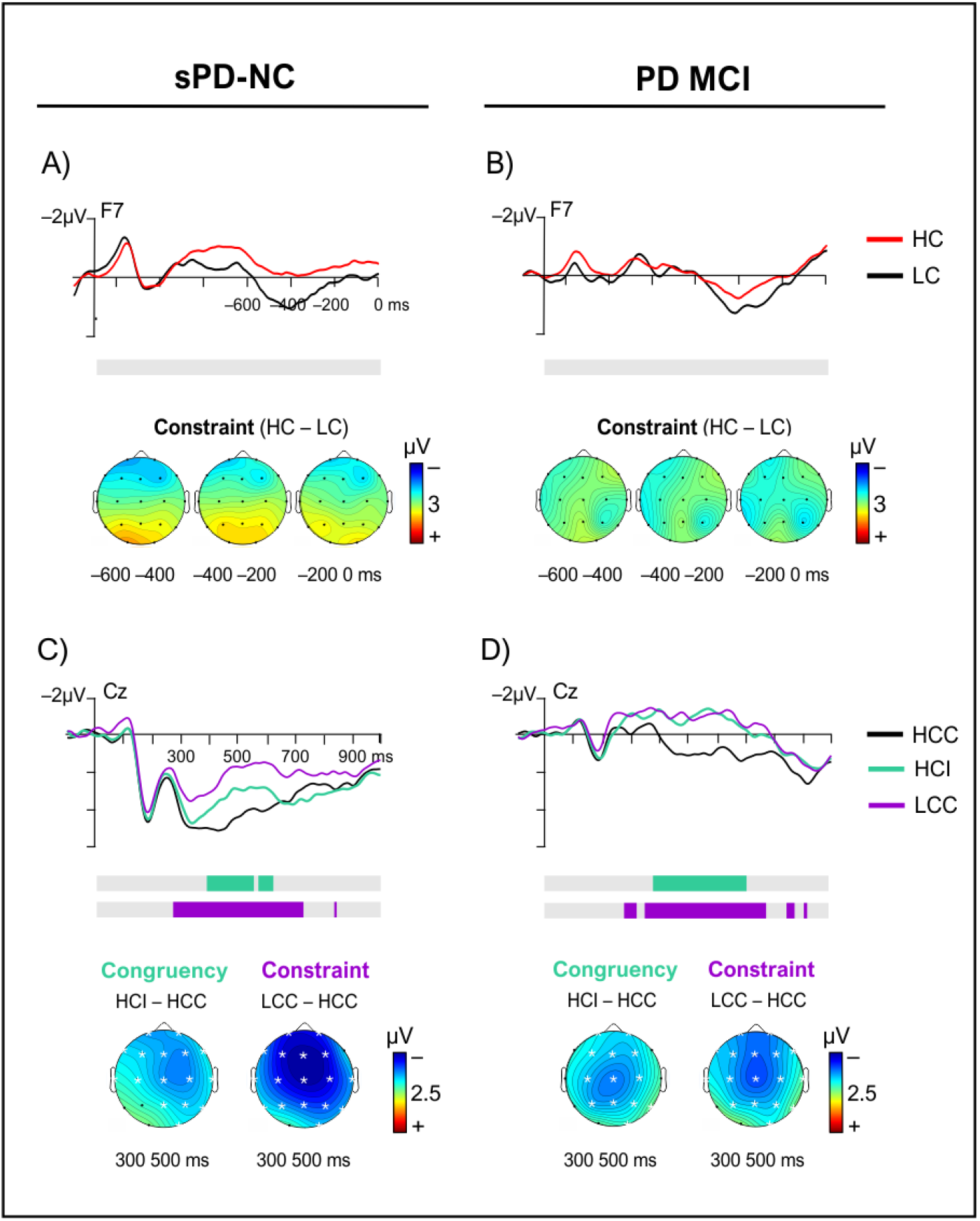
ERP results for sPD-NC and PD-MCI groups. **A and B)** Grand-averaged ERPs at the F7 electrode during the pre-word interval for (A) sPD-NC and (B) PD-MCI. ERPs are time-locked to the presentation of the penultimate word (100 pre-stimulus baseline). The horizontal bar highlights (grey) the time interval in which significant differences were obtained. Below the ERPs, topographical maps indicating the electrodes (white asterisks) that showed a significant effect of constraint (HC minus LC) in the 600 ms prior to the critical word (in steps of 200 ms). **C and D)** Grand-averaged ERPs at Cz electrode after the presentation of the sentence-final word (100 pre-stimulus baseline) for (C) sPD-NC and (D) PD-MCI. The horizontal bars highlight the time interval of the N400 effect of Congruency (green) and Constraint (purple). Topographical maps with electrodes (white asterisks) that showed effects in the time interval in which N400 effects were expected to be maximal (300 to 500 ms post-word onset).

After the presentation of the final word (**Fig. 3C** and **Fig. 3D**) there was an observable larger negativity for unexpected (HCI and LCC) relative to expected words (HCC) in both groups, in line with classical N400 context effects. These observable differences were confirmed by significant clusters as specified next.

There were significant effects of constraint (HCC versus LCC) both for the group of patients without MCI (p <.001) (**Fig. 3C**) and the group with MCI (p <.001) (**Fig. 3D**). That is, as expected, words in less predictive contexts elicited a larger N400 than words in strongly predictive contexts, a difference that encompassed the interval between about 300 to 700 ms post word onset. The scalp distribution of the effect covered a wide set of electrodes over frontal and central regions of the scalp, with a slight frontward amplitude maximum. No differences between groups were found.

As for the effect of congruency in predictive contexts (HCC versus HCI), again as expected, significant clusters reflected that incongruent words had more negative amplitudes than congruent words in patients without MCI (p = .005) (**Fig. 3C**), and also in with MCI (p = .001) (**Fig. 3D**). Relevantly, in this case, there were significant differences between the groups (**Fig. 4**). The difference waveforms (HCI minus HCC) of each group were contrasted in a between-group cluster-based permutation test including all electrodes and time-points (from 0 to 1000 ms). The test yielded a significant cluster (p = .008) that revealed a longer-lasting effect of congruency for the group of patients with MCI, which can be visualized in **Figure 4A**. More specifically, the cluster spanned approximately from 630 to 760 ms after word onset at centro-parietal sites (significant electrodes: C3, Cz, C4, P3, Pz, P4, T5, T4, T6, O2) (see the scalp distribution of *t*-values in **Fig. 4B**). This result suggests a prolonged N400 congruity effect in the PD-MCI group relative to the group of patients without MCI.

**Figure 4.**
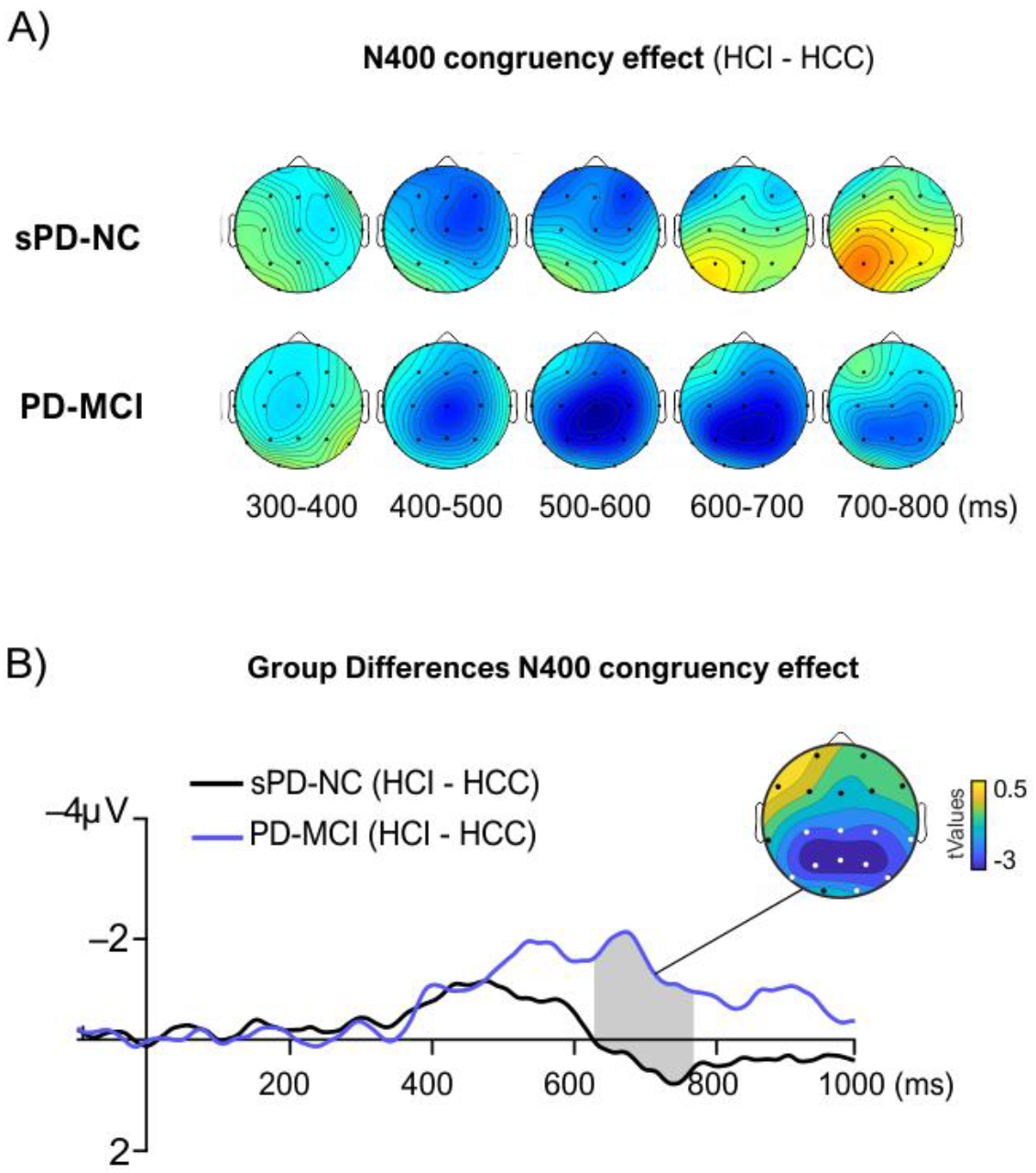
**A)** Topographical maps showing the temporal evolution and scalp distribution of the congruency contrast (HCI minus HCC) during final word processing, showing a prolonged N400 congruency effect in the group of individuals with PD and MCI. **B**) Grand-averaged ERPs of the difference waveforms (HCI and HCC) of both groups, averaged over the set of electrodes that were part of the significant cluster of differences between the groups (highlighted in white in the topographical map). The grey-colored area shows the timepoints included in the cluster. The topographical map shows the scalp distribution of the averaged t-values within the cluster (cluster t-value ± 2.02 for an alpha level of .05).

### 3.2. Correlations between ERP effects and verbal fluency scores

After establishing the ERP effects, we examined the association between ERP measures and verbal fluency scores in all PD patients (N = 78) (for further specification of this analysis, see **Section 3.2**) by means of non-parametric Spearman correlations between direct scores in verbal fluency tests, and four ERP measures, namely, 1) SNP (LC minus HC), 2) N400 constraint effect (LCC minus HCC), 3) N400 congruency effect (HCC minus HCI), and 4) prolonged N400 congruency effect (HCC minus HCI). Note that, in these analyses, the direction of the condition subtraction is reversed, such that positive values represent differences in the expected direction of the effect (e.g., in the case of the SNP, positive values indicate larger amplitudes for HC than LC). All correlations were corrected using Holm-Bonferroni correction (indicated as pHB in the text).

The results of the correlational analyses are presented in **Table 3**. After correcting for multiple comparisons, two of the ERP measures showed a significant correlation with verbal fluency scores: the SNP and the prolonged N400 congruity effect (**Fig. 5**). In particular, the SNP showed a significant positive correlation with verbal fluency (r(77) = .326, pHB = .010). Note that we excluded one participant who had outlying scores in the SNP (z-score = −3.91) although the correlation was unaffected by its inclusion (r(78) = .326, pHB = .01), which is normal when using nonparametric Spearman correlations that are very robust to the presence of outliers. This indicates that a larger SNP (more negative amplitudes prior to words in predictive than in unpredictive contexts) is associated with better verbal fluency performance in patients with PD. In turn, the prolongation of the N400 congruency effect (more negative amplitudes for incongruent than congruent words 600-800 ms post-word) correlates with worse scores in verbal fluency (r(78) = – .358, pHB = .005). No other correlations were statistically significant.

**Table 3.**
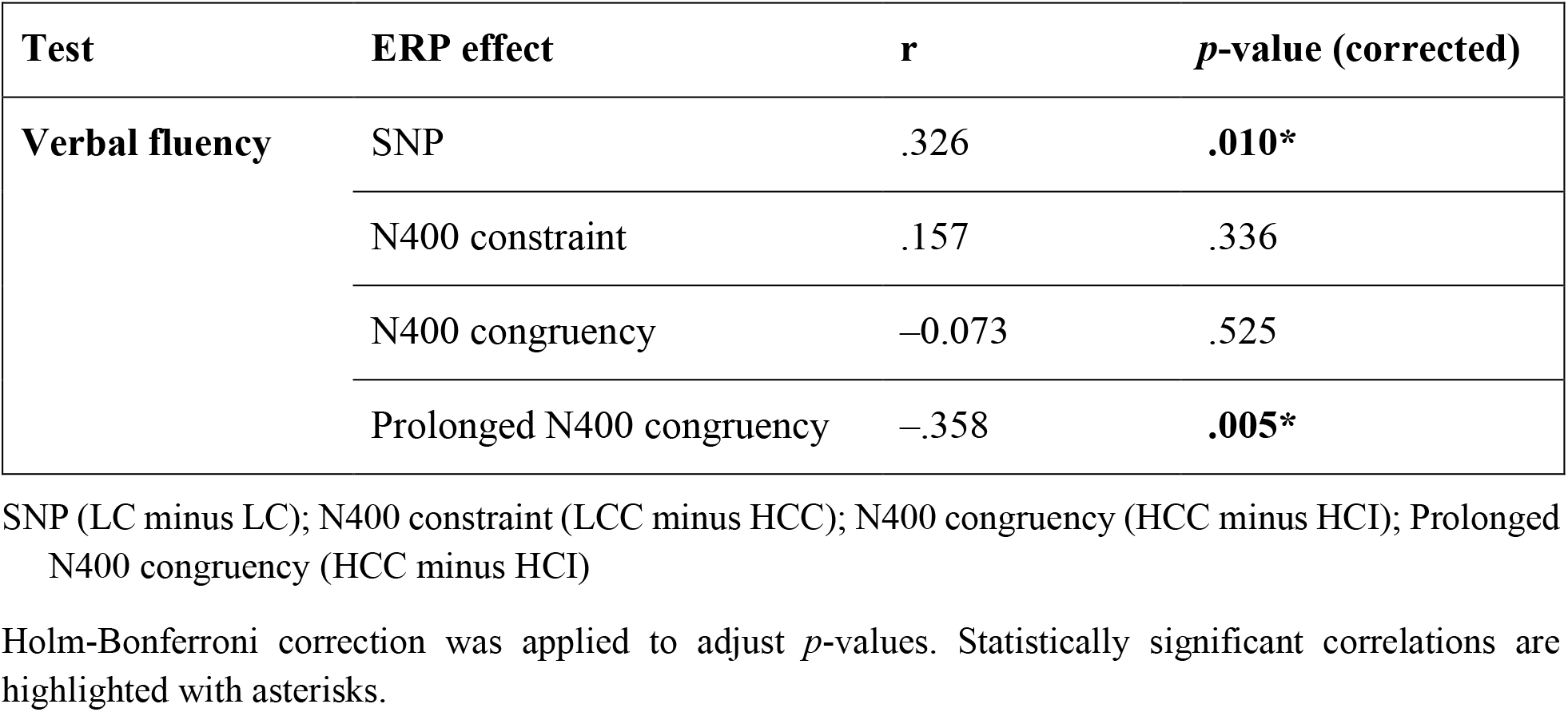
Results of correlational analyses between ERP measures and verbal fluency scores.

**Figure 5.**
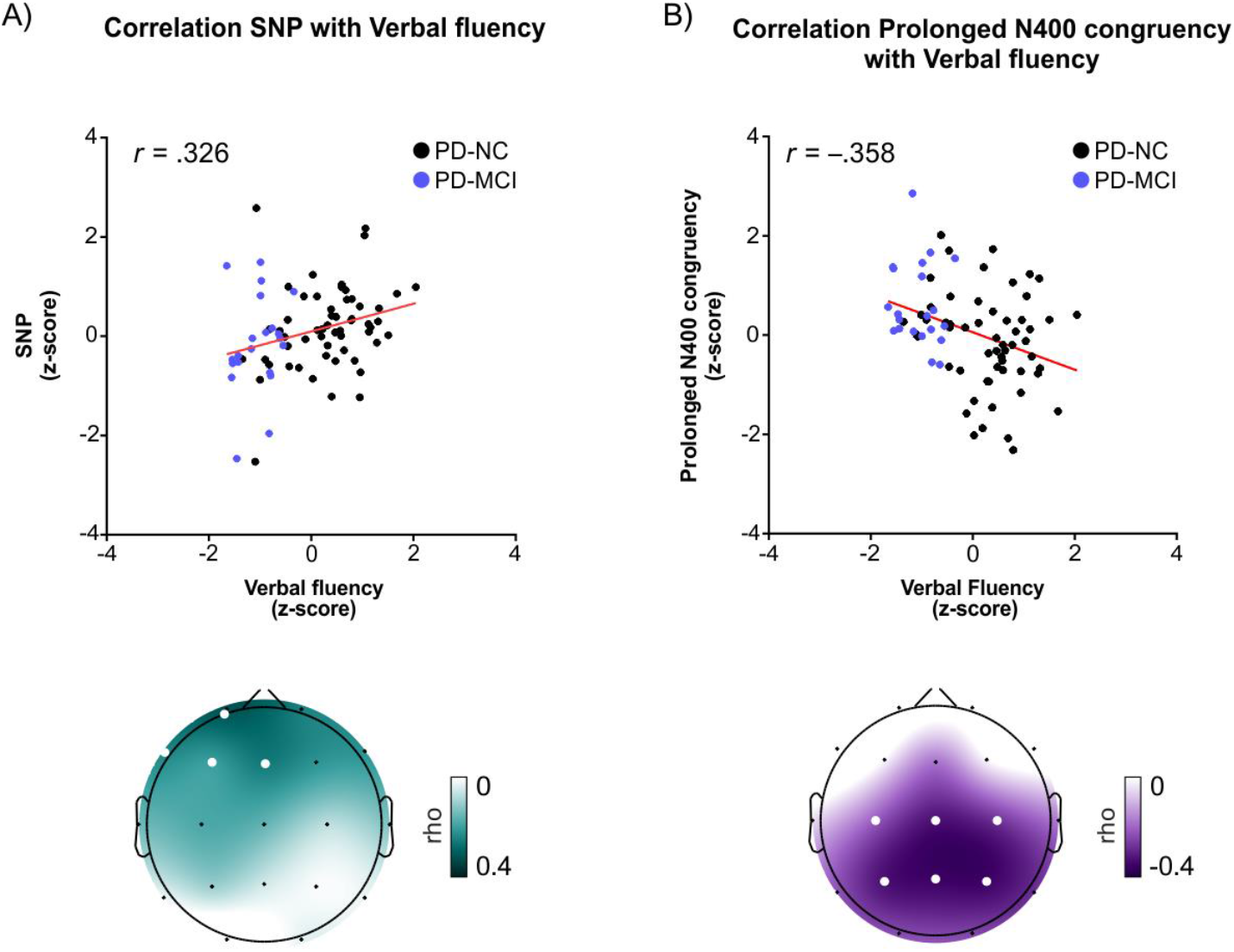
Scatterplots and scalp maps of significant correlations between ERP measures and verbal fluency scores in all PD patients (PD-NC and PD-MCI) (N = 78). **A)** Scatterplot of the positive correlation (r = .326, pHB = .010) between verbal fluency scores and the SNP (more negative amplitudes prior to words in HC than in LC contexts) and **B**) of the negative correlation (r = –.358, pHB = .005) between verbal fluency scores and the prolongation of the N400 congruency effect (more negative amplitudes for incongruent than congruent words 600-800 ms post-word). All values were normalized. Below the scatter diagrams, the r-values (Spearman rho) for all 19 electrodes are plotted on an idealized head. Darker shading indicates larger r-values in the direction of the correlation. Electrodes included in the ROI used to perform the reported correlations with verbal fluency (shown in the scatter diagrams) are highlighted (in white). R-values between electrodes were estimated by spherical spline interpolation.

## 4. DISCUSSION

Motivated by previous evidence of altered predictive language processing in populations with low cognitive resources (e.g., Federmeier *et al*., 2002; 2005; 2010), the present study assessed language prediction in PD with normal cognition (PD-NC) and PD with MCI (PD-MCI). To this end, ERP modulations were examined before and after encountering words that could or could not be predicted from context, thus capturing both the anticipatory and processing stage of prediction. Specifically, we focused on SNPs as signatures of semantic anticipation, and post-word N400 context effects as indices of predictive pre-activation. Relative to controls, PD-NC patients exhibited expected ERP signatures, pointing to preserved predictive language processing. In turn, PD-MCI patients showed absent SNPs and a significant prolongation of N400 congruency effects in predictive contexts, suggesting altered mechanisms tied to language prediction. Interestingly, correlational analyses revealed that worse verbal fluency performance was associated with the presence of the ERP pattern observed in the PD-MCI group.

### Neural correlates of predictive sentence processing are preserved in PD-NC

Consistent with prediction, PD-NC patients exhibited the canonical reduction of N400 amplitude for expected (HCC) compared to unexpected words (LCC and HCI) (Kutas *et al*., 1984). This effect is taken to reflect facilitated lexical and semantic activation of expected words as a result of their pre-activation (Kutas *et al*., 1999; Lau *et al*., 2008). In line with this interpretation, in predictive contexts (HC), PD-NC individuals exhibited a frontally distributed negative SNP that developed progressively prior to the presentation of the final word (León-Cabrera *et al*., 2017; 2019; for similar results, see Grisoni *et al*., 2017; Li *et al*., 2017). Studies in healthy population suggest that such SNPs capture anticipation of semantic aspects of the upcoming word (for a recent review, see Pullvermüller *et al*., 2020). More specifically, it may reflect general-domain of language-specific mechanisms supporting retrieval or maintenance of pre-activated representations (Li *et al*., 2017; León-Cabrera *et al*., 2019). Altogether, PD-NC patients seem to make proper use of sentence contexts, affording normal semantic processing in PD, as has been previously observed (Friederici *et al*., 2002). In addition, the current study shows, for the first time, that they can also use sentence contexts to anticipate and pre-activate upcoming information.

Previous N400 studies have pointed to altered lexical and semantic activation in PD-NC (Angwin *et al*., 2017; Kutas *et al*., 2013; Angwin *et al*., 2004; Arnott *et al*., 2010, Copland *et al*., 2009; Angwin *et al*., 2007). However, these studies employed single-word contexts, instead of sentence contexts. Importantly, sentence contexts involve the construction of a high-level meaning representation (Graesser *et al*., 1994), which provides additional constraints to pin down relevant lexical and semantic representations, perhaps compensating for potential activation problems within semantic networks. Nonetheless, note that all patients in the current study were on dopaminergic medication. PD-NC patients on dopaminergic medication perform better in semantic priming tasks than PD-NC patients off medication (Angwin *et al*., 2004a; Angwin *et al*., 2009), suggesting that dopamine (DA) mediates semantic activation, arguably through calibration of the spread and focus of activation within semantic networks (Kischka *et al*., 1996). DA has also been linked with anticipatory processes in PD. Specifically, PD-NC patients off medication show reduced amplitudes of the stimulus-preceding negativity (SPN), suggesting that the DA system is implicated in the anticipation of motivationally salient and rewarding stimuli (Mattox *et al*., 2006). Future studies testing patients off medication would help untangle to what extent dopaminergic compensation contributed to the preservation of otherwise altered mechanisms.

### PD-MCI impacts late semantic processing in predictive contexts

To further investigate the impact of cognitive impairment, we also examined neural responses in a smaller sample of PD-MCI patients. Compared to matched PD-NC patients (sPD-NC), PD-MCI patients exhibited expected N400 amplitude modulations within the normal onset latency (i.e., about 300-400 ms post-word onset). However, most remarkably, PD-MCI patients showed a significantly prolonged N400 congruency effect in predictive contexts, extending up to 700 ms after word onset. Critically, this was not observed in the N400 constraint contrast (LCC versus HCC), which suggests that the prolongation stemmed from the ERP response to incongruent words, rather than to congruent words (i.e. from HCI, instead of HCC). With that in mind, it is important to note that N400 context effects do not solely capture prediction effects, but also integration demands (van Berkum *et al*., 1999). Interestingly, a recent study showed that prediction effects dominate the earlier portion of the N400 context effect (starting as early as 200 post-word onset), whereas integration effects began later and continued until about 650 ms after word onset (Nieuwland *et al*., 2020; see also, Brothers *et al*., 2014; Lau *et al*., 2016). As such, prolonged N400 congruency effects may reflect abnormally sustained efforts to integrate the incongruent word with the previous context (Nieuwland *et al*., 2020). Curiously, a previous study found that patients with bilateral basal ganglia lesions exhibited similarly prolonged N400 effects in sentence processing (up to 700 ms post-word) (Kotz *et al*., 2003). In PD, the worsening of the condition from PD to PD-MCI may lead to altered late integrational semantic processing as well.

More recently, N400 effects have postulated to reflect the ‘prediction error’ (albeit with nuances in their definition, see Willems *et al*., 2016; Rabovsky *et al*., 2018; Kuperberg *et al*., 2020). In predictive processing frameworks (Clark, 2013), predictions are sent top-down from higher to lower level regions to ‘explain away’ the incoming sensory input and, then, the ‘prediction error’ – the portion of information that remains unexplained – is sent back and used as a learning signal to update internal models and improve future predictions. Thus, prolonged N400 congruency effects may indicate difficulties in this ‘learning’ phase, leading to less efficient internal model updating. In line with less efficient predictive language processing in PD-MCI, there was no SNP in this group, pointing to absent semantic anticipation. In this case, however, the effect was absent in the sPD-NC group as well, and therefore cannot be attributed to MCI.

### Worse verbal fluency performance is associated with altered signatures of predictive language processing in PD

Interestingly, correlational analyses revealed that lower verbal fluency scores were associated with reduced SNPs and prolonged N400 congruency effects. Similarly, previous evidence has shown that verbal fluency scores are a good predictor of the status of predictive language processing in healthy older population (Federmeier *et al*., 2002). Worse performance in verbal fluency is generally associated with executive dysfunction, such as difficulties in rule switching or inhibition. Executive dysfunction is common with the progression of fronto-striatal deterioration in PD and in PD-MCI (Kudlicka *et al*., 2011; Monchi *et al*., 2004; Aarsland *et al*., 2010) and has been proposed to indirectly hinder language processing in PD (Grossman *et al*., 2002; 2003). However, verbal fluency performance depends not only on frontal lobe function, but also on lexical and semantic retrieval processes dependent on temporo-parietal structures (Unsworth *et al*., 2011). In fact, recent findings point to a greater weight of language-related processes than executive function in verbal fluency (Whiteside *et al*., 2016). Recent meta-analyses and lesion-based studies have also shown that verbal fluency is supported by standard language networks and underlying white-matter connectivity (Griffis *et al*., 2017; Costafreda *et al*., 2006; Baldo *et al*., 2006). Therefore, the ERP pattern associated with lower verbal fluency – suggestive of less efficient predictive processing mechanisms, as previously dicussed – may reflect not only executive dysfunction, but also disruption in semantic networks, in line with recent findings of difficulties in temporal-dependent functions in PD and PD-MCI (Horta-Barba *et al*., 2020; Martínez-Horta *et al*., 2019; Lang *et al*., 2019).

### Concluding remarks and future directions

Overall, the results suggest preserved predictive language processing in PD-NC. In turn, in PD-MCI, further cognitive limitations hinder mechanisms associated with semantic prediction in normal circumstances. While these limitations may not prevent the pre-activation of relevant representations, they might negatively affect sentence processing due to poorer semantic anticipation and less efficient semantic processing when contexts are predictive. Furthermore, both executive dysfunction and damage within semantic networks may underlie these difficulties, as suggested by their association with low verbal fluency scores. Along with recent research (Horta-Barba *et al*., 2020; Martínez-Horta *et al*., 2019; Lang *et al*., 2019), these findings emphasize the value of examining cognitive changes in PD and PD-MCI in domains beyond executive functions, like the language domain. Interestingly, altered N400 effects are associated with a higher risk of transiting to dementia in individuals with amnesic MCI (Olichney *et al*., 2008). Therefore, future longitudinal studies in PD could evaluate changes in N400 congruency effects as potential markers of the transition to PD-MCI and PDD (for interested readers, see Supplementary Materials for a hierarchical regression analysis further attesting that the N400 congruency effect is a good predictor of global cognitive decline in PD). Finally, as previously highlighted, future studies testing patients off medication could assess the exact role of DA in predictive sentence comprehension. Yet, the value of evaluating PD patients on stable doses of dopaminergic medication must be emphasized, as it is representative of their habitual cognitive status and thus has the potential to uncover limitations that may impact the everyday functioning of individuals with PD.

## ACKNOWLEDGMENTS

In honor and loving memory of Dr. Jordi Riba, who passed away in the process of preparing this manuscript. We would like to thank Dr. Ion Iarritu, Dr. Laura Ferreri, Miriam Cornellà and Fernando Martínez for their help with data collection and technical support.

## FUNDING

This study was funded by grants from the Fundació la Marató de TV3 2014/U/477 and 20142910, and by financial support from the Center for Biomedical Research and Neurodegenerative Resources (CIBERNED). PLC was funded with a pre-doctoral grant FPU15/05554 (FPU “Ayudas para la Formación de Profesorado Universitario”) of the Spanish Ministry of Education, Culture and Sport. JP was funded by PERIS (expedient number: SLT008/18/00088) of Generalitat de Catalunya. JK and HB-K were funded by FIS PI18/01717. HB - K was funded by Río Hortega CM17/00209 of Instituto de Salud Carlos III (ISCIII) (Spain). JM was funded by Río Hortega CM15/00071 of Instituto de Salud Carlos III (ISCIII) (Spain). None of these funding sources had any involvement in the conduct of the research or the preparation of the article.

## COMPETING INTERESTS

The authors declare no disclosure of financial interests and potential conflict of interest.

## Notes

### Competing Interest Statement

The authors have declared no competing interest.

