## Supplementary Materials for "Neural signatures of predictive language processing in parkinson’s disease with and without mild cognitive impairment"

### 1. Correlational analyses between ERP measures and verbal fluency (semantic and phonological separately)

To provide a more complete picture of the relation between the ERP measures and verbal fluency performance, we additionally performed a correlational analysis with semantic and phonological fluency separately. The parameters used to compute the ERP measures were the same as those used for the analyses in the main paper (see Section 3.2 in the Methods). For each group, the mean and standard deviation of the scores in semantic and phonological fluency tests are presented in **Table** 1.

**Table 1. Demographics and clinical features of the patient and control samples.**

| | **CG** | **PD-NC** | P | **sPD-NC** | **PD-MCI** | P |
|---|---|---|---|---|---|---|
| N | 24 | 58 | | 19 | 20 | |
| Semantic fluency (DS) | 22.08 (3.67) | 20.03 (5.10) | .07 | 19.63 (6.29) | 12.05 (3.57) | <.001 |
| Phonological fluency (DS) | 17.25 (3.99) | 16 (4.9) | .27 | 15.63 (4.16) | 9 (3.22) | <.001 |

The results of the correlational analysis are shown in **Table 2.** Two correlations survived after correcting for multiple comparisons (using Holm-Bonferroni correction, indicated as pHB), which were essentially the same as in the main paper, but only with phonological fluency in this case. Specifically, phonological fluency scores correlated positively with the SNP (r(78) = .393, pH = .003) and a negatively with prolonged N400 effects (r(78) =–371, pHB = .005) (see **Figure** 1). Note, however, that the correlations between these ERP measures (SNP and prolonged N400 congruency effects) and semantic fluency scores were significant or marginally-significant before correcting (p = 056 with the SNP; and p = .01 with the N400 congruency effect). Indeed, the correlation with the N400 congruency effect was marginally non-significant even after correction (p = .063). Likewise, the correlation between semantic fluency and the N400 constraint effect remained marginally non-significant after correction (p = .063).

**Table 2. Results of correlational analyses between ERP measures and verbal fluency scores (semantic and phonological fluency)**.

| Test | ERP effect | r | *p*-value | Test | ERP effect | r | *p*-value |
|---|---|---|---|---|---|---|---|
| **Semantic fluency** | SNP | .217 | .056 (.225) | **Phonological fluency** | SNP | .393 | **<.001* (.003*)** |
| | N400 constraint | .287 | **.010*** (.063) | | N400 constraint | –.020 | .856 (1.713) |
| | N400 congruency | -.007 | .949 (1.713) | | N400 congruency | –.118 | .300 (.902) |
| | Prolonged N400 congruency | -.284 | **.011*** (.063) | | Prolonged N400 congruency | –.368 | **<.001* (.006*)** |

SNP (LC minus LC); N400 constraint (LCC minus HCC); N400 congruency (HCC minus HCI); Prolonged N400 congruency (HCC minus HCI)

Corrected p-values (Holm-Bonferroni correction) are shown in brackets. Statistically significant correlations are highlighted with asterisks.

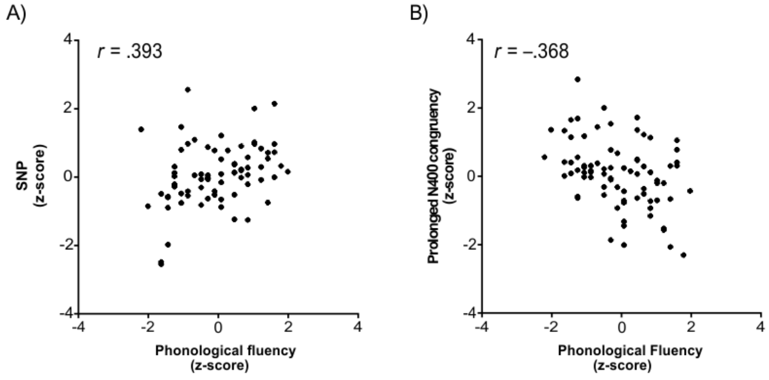


**Figure** 1**. Scatterplots of significant correlations between ERP measures and phonological verbal fluency in all PD patients (with and without MCI). A)** Scatterplot showing the significant positive correlation between phonological fluency scores and the SNP (more negative amplitudes prior to words in predictive than in unpredictive contexts). All values were normalized. **B**) Scatterplot of the significant negative correlation between phonological fluency scores and a prolongation of the N400 congruency effect (more negative amplitudes for incongruent than congruent words 600-800 ms post-word). All values were normalized.

### 2. Linear regression between PDCRS scores and prolonged N400 congruency effects

Based on the finding of a prolonged N400 congruency effect, we further investigated if this effect was a good predictor of global cognitive function, operationalized as the PD-CRS total score, in the whole sample of individuals with PD (N = 78), controlling for age and education.

For each participant, we quantified the prolonged N400 congruency effect (hereafter referred to as Late N400) as the mean amplitude difference between HCI and HCC in all electrodes that showed a significant difference in the exploratory analyses: C3, Cz, C4, P3, Pz, P4, T5, T4, T6, O2) in the 600 to 800 ms interval (time-locked to final-word onset). Note that the electrodes used to compute the prolonged N400 effect here are different from those used in the main text (C3, Cz, C4, P3, Pz, P4). This is because the current analysis was tailored for the prolonged N400 congruency effect exclusively, whereas, in the main text, we used the same electrodes for all N400 effects (i.e., congruency and constraint) for parsimony and comparability.

A two-stage hierarchical multiple regression model was performed to predict the PD-CRS using age and education at the first stage and adding the prolonged N400 congruency effect at the second stage (**Table 3**). This procedure allowed to evaluate the unique contribution of the ERP measure to explain global cognitive performance while controlling for age and education. The first model accounted for 27% of the variance ($R^2$ = .275; F(2,77) = 14.24, $p$ < .001) and both age and education were significant predictors of the PDCRS total score (both $p$ < .03). The addition of the Prolonged N400 measure measure as a predictor significantly improved the model ($F_{change}$(2,75) = 10.25, $p$ = .002), explaining 36 % of the PD-CRS score ($R^2$ = .363; F(3,74) = 14.08, $p$ < .001) (**see Fig. 2**). All predictors were significant in the second and final model (all $p$-values < .03). All assumptions for multiple linear regression were met. There were no problems of multicollinearity (all tolerance values were higher than .2 and the variance inflation factor was higher than 10) (Menard, 1995; Myers, 1990), or outliers or influential cases (all cases had Cook's distance > 1 and Mahalannobis values > 15) and residuals were equally and normally distributed (assessed through plot inspection).

**Table 3.** Coefficients for the two-stage hierarchical multiple regression with PDCRS as a dependent variable and age, education (stage 1) or age, education and prolonged N400 congruency effect (stage 2) as predictors.

| | **Unstandardized coefficients** | | **Standardized coefficients** | | |
|---|---|---|---|---|---|
| **Model 1** | **B** | **SE** | **β** | **t** | **p** |
| (Constant) | 135.33 | 18.49 | | 7.31 | < .001 |
| Age | –.830 | .241 | –.370 | –3.45 | .001 |
| Education | .897 | .382 | .252 | 2.35 | .021 |
| **Model 2** | | | | | |
| (Constant) | 125.52 | 17.71 | | 7.08 | <.001 |
| Age | –.652 | .234 | –.291 | –2.78 | .007 |
| Education | .850 | .361 | .239 | 2.35 | .021 |
| Prolonged N400 congruency effect | 2.2 | .693 | .309 | 3.20 | .002 |

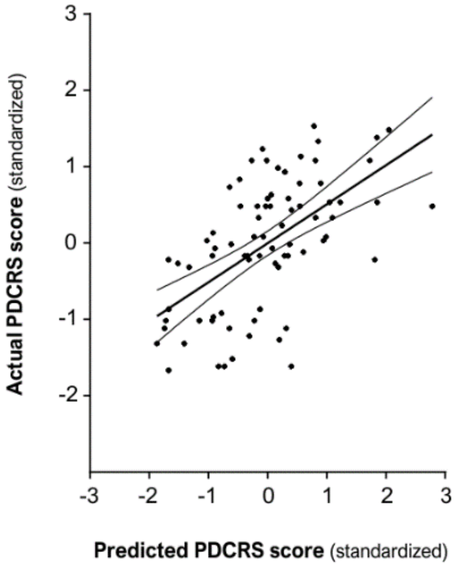


**Figure 2.** Scatterplot of observed PDCRS scores (Y axis) by predicted PDCRS scores (X axis) obtained in the linear regression model with age, education and the Prolonged N400 congruency effect (HCI minus HCC difference waveform in the 600-700 ms over a centro-parietal cluster), capturing the prolongation of the N400 congruity effect, as predictors. Note that smaller values of the Prolonged N400 congruency effect correspond to larger N400 effects (i.e. the subtraction between HCI and HCC yielding a negative value, indicating more negative amplitudes for incongruent than congruent words). Scores are standardized. A linear regression line was fitted to the data with 95% confidence intervals.

**References (Supplementary Materials)**

Myers, R. H. (1990). Detecting and combating multicollinearity. Classical and modern regression with applications, 368-423.

Menard, S. (1995) Applied Logistic Regression Analysis: Sage University Series on Quantitative Applications in the Social Sciences, Thousand Oaks, CA: Sage.
